# Methylglyoxal alters C-fibre Activity-Dependent Slowing and Induces Heat Hyperalgesia in a Sex-Dependent Manner

**DOI:** 10.1101/2025.10.09.680671

**Authors:** Atanaska N Velichkova, Menekse Mutlu-Smith, Amy L Hall, Carole Torsney

## Abstract

**Introduction:** Raised plasma levels of the glycolytic metabolite methylglyoxal are associated with pain symptoms in patients with diabetic neuropathy. Methylglyoxal can regulate the function of Na_V_1.7 and Na_V_1.8 ion channels that are involved in the phenomenon of activity-dependent slowing (ADS) in C-fibre nociceptors. C-fibre ADS differs between the sexes and can regulate spinal network function.

**Objective:** Explore the impact of methylglyoxal upon C-fibre ADS and pain sensitivity in both sexes. In addition, investigate the influence of ADS upon the processing of C-fibre inputs, in noxious heat responsive spinal neurons.

**Results:** Compound action potential recording of isolated dorsal roots incubated with methylglyoxal (100µM, 3 hr) revealed a sex-dependent impact upon C-fibre ADS. In male roots, C-fibre ADS was reduced whereas in female roots it was increased. Acute methylglyoxal application (100µM/1mM, 10 min) did not modify C-fibre ADS. Systemic methylglyoxal administration (5μg, 3h prior) induced heat hyperalgesia in male but not female juvenile rats. Patch-clamp recording in spinal slices with attached dorsal roots revealed that length-dependent manipulation of ADS altered action potential firing to stimulus trains in noxious heat responsive Fos-EGFP+ spinal neurons.

**Conclusion:** We propose that methylglyoxal sex-dependent regulation of C-fibre ADS influences the spinal processing of noxious heat inputs that may contribute to the male specific induction of heat hyperalgesia following systemic methylglyoxal treatment.

**Summary:** Methylglyoxal, implicated in painful diabetic neuropathy, sex-dependently alters C-fibre ADS, potentially influencing spinal processing of noxious heat to drive heat hyperalgesia in males only.

## Introduction

Diabetes is a significant global health challenge, affecting 11% of the adult population, with projections that 1 in 8 adults will be living with diabetes by 2050 (International Diabetes Federation). Up to half of diabetes patients develop neuropathy[47] with 20-40% of diabetes patients suffering from painful diabetic neuropathy[31]. The highly reactive glucose-derived metabolite methylglyoxal (MG) is a risk factor for diabetic neuropathy[3] and raised plasma levels discriminate diabetic neuropathy patients with painful symptoms compared to those without[10]. Post-translational modification of Na_V_1.8 in sensory neurons is implicated in the pain promoting action of MG[10].

Na_V_1.8 and Na_V_1.7, whose function is also affected by MG[10], are involved in activity-dependent slowing (ADS) in C-fibre nociceptors[5; 29; 30; 45; 57], with the ratio of Na_V_1.7 to Na_V_1.8 influencing the extent of ADS[45]. ADS manifests as a progressive slowing of action potential conduction velocity in response to repeated stimulation[22; 52; 56; 62], thus regulating intervals between successive action potentials being relayed along nociceptor axons, influencing the temporal delivery to and processing of pain in the CNS[16]. It is frequency- and length-dependent, with greater ADS at higher frequencies[22; 52; 56; 62] and over longer lengths[16; 49; 64]. C-fibre ADS is altered in chronic pain patients[32-34; 42; 53] including those with diabetic neuropathy[33]. It is also altered in preclinical pain models[16; 20; 21; 41; 54; 60; 61] including diabetic neuropathy models[21; 61]. Specifically, we have shown that C-fibre ADS and its alteration in preclinical models is sex-dependent[16; 60], may relate to sex differences in heat pain and likely involves sex differences in the Na_V_1.7/ Na_V_1.8 ratio[60].

The aim of this study was therefore to explore the impact of MG upon C-fibre ADS and heat pain sensitivity in both sexes. In addition, the impact of ADS upon spinal processing of noxious heat inputs was investigated.

## Methods

### Animals

All experiments were approved by the University of Edinburgh Ethical Review Committee and carried out in accordance with the UK Animal (Scientific Procedures) Act 1986. Juvenile Sprague–Dawley (SD) rats, aged approximately postnatal Day 21 (∼P21) (University of Edinburgh Bioresearch and Veterinary Services) and adult wild type (WT) *C57BL/6* and hemizygous Fos-EGFP mice (B6.Cg-Tg(Fos/EGFP)1-3Brth/J, Jackson), aged 5-11 weeks, of both sexes were used. Animals were housed in cages at 21°C and 55% relative humidity, with a 12-h light–dark cycle and food and water provided *ad libitum*.

### Methylglyoxal-induced model of diabetic behavioural hypersensitivity in juvenile rats

SD rats of both sexes received 0.2 ml intraperitoneal (i.p.) injection (without anaesthesia) of methylglyoxal (MG) (5μg) or vehicle (sterile 0.9% w/v saline) solution 3h prior to sensory testing.

### Behavioural sensory testing in juvenile rats

Sensory testing was performed as described previously [16; 60]. Briefly, after habituation on an elevated glass platform maintained at 30°C, a radiant heat stimulus (Hargreaves apparatus, IITC Life Science, CA, USA) was applied to the midplantar surface of the hindpaw (X3 to calculate the average) to determine the thermal withdrawal latency of the nociceptive flexion withdrawal reflex. Following habituation on an elevated mesh platform, the mechanical threshold of the nociceptive flexion withdrawal reflex was determined using an electronic von Frey apparatus (Ugo Basile), applied to the midplantar surface of the hindpaw (X3 to calculate average). Animals were tested before and 3 hours following i.p MG/vehicle administration.

### Activation of noxious heat responsive spinal neurons in Fos-EGFP mice

To selectively target noxious heat responsive neurons for *ex vivo* electrophysiological investigation hemizygous Fos-EGFP mice were used. *In vivo* intraplantar capsaicin injection or natural noxious heat stimulation, were used to selectively activate noxious heat responsive spinal neurons.

Capsaicin injections were administered subcutaneously into the plantar surface of the left hindpaw (0.25mg/ml, 10μL), under isoflurane anaesthesia. Natural noxious heat stimulation was applied by immersing one hindpaw in a 52°C water bath, under isoflurane anaesthesia, twice for 15 secs with a 2 minute resting interval, as conducted previously[1]. Approximately 2 hours post-stimulation, when Fos-EGFP expression is maximal[8], animals were terminated for preparation of *ex vivo* spinal cord slices.

### Isolated dorsal root and spinal cord slice preparations

Isolated dorsal roots and spinal cord slices with attached dorsal roots were prepared as described previously[16; 58; 60]. Under isoflurane-induced anaesthesia, naive rats (∼P21) and adult WT *C57BL/6* and hemizygous Fos-EGFP mice (5-11 weeks) were decapitated and spinal cords, with attached dorsal roots, were removed in an ice-cold dissection solution, containing (in mM) 3.0 KCl, 1.2 NaH_2_PO_4_, 26 NaHCO_3_, 15 glucose, 251.6 sucrose, 7 MgCl_2_, and 0.5 CaCl_2_, pH 7.3–7.4. For dorsal root preparations, the lumbar (L4/L5) dorsal roots were cut near the entry zone, and dorsal root ganglia were removed. For spinal cord slice preparations, the lumbar (L4/L5) segment with attached dorsal roots was embedded in an agarose block, and 450-500 μm slices were cut. The tissue was briefly recovered (up to 15 mins) in 32-34°C oxygenated *N*-methyl-D-glucamine (NMDG) recovery solution, containing (in mM) 93 NMDG, 2.5 KCl, 1.2 NaH_2_PO_4_, 30 NaHCO_3_, 25 glucose, 20 4-(2-hydroxyethyl) piperazine-1-ethanesulphonic acid (HEPES), 5 sodium absorbate, 2 thiourea, 3 sodium pyruvate, 10 MgSO_4_, and 0.5 CaCl_2_, pH 7.3–7.4, and then incubated at room temperature in oxygenated holding solution, comprising (in mM) 92 NaCl, 2.5 KCl, 1.2 NaH_2_PO_4_, 30 NaHCO_3_, 25 glucose, 20 HEPES, 5 sodium absorbate, 2 thiourea, 3 sodium pyruvate, 2 MgSO_4_, and 2 CaCl_2_, pH 7.3–7.4, for a minimum of 1h. Electrophysiological recordings were conducted in continuous perfusion with recording solution containing (in mM) 125.8 NaCl, 3.0 KCl, 1.2 NaH_2_PO_4_, 26 NaHCO_3_, 15 glucose, 1.3 MgCl_2_, and 2.4 CaCl_2_, pH 7.3–7.4.

### Dorsal root chronic methylglyoxal treatment

Dorsal roots were recovered in oxygenated holding solution with 100μM MG or vehicle (distilled water) for 3 hours prior to recordings, which were conducted in the continuous presence of recording solution with 100μM MG or vehicle.

### Compound action potential (CAP) recording of C-fibre activity-dependent slowing

CAPs were recorded using two glass suction electrodes placed at each end of the dorsal root, one for electrical stimulation (ISO-Flex stimulus isolator, AMPI, Jerusalem, Israel) and the other for recording. The C-fibre activation threshold was defined and the response amplitude and conduction velocity, measured at 500 μA, as described previously[16; 60]. Data were acquired and recorded using an ER-1 differential amplifier (Cygnus Technologies, PA, USA) and pCLAMP™ 10 software (Molecular Devices, Wokingham, UK). Data were filtered at 10 kHz and sampled at 50 kHz.

To assess C-fibre ADS, repetitive stimulation (40x stimuli) was applied at C-fibre intensity (500 μA intensity; 0.1 ms duration) at 1, 2 and 10 Hz with 10 min recovery intervals between successive repetitive stimulations. To investigate C-fibre ADS following acute (10 minute) MG treatment, 2 Hz stimulation was applied following 100μM and 1mM MG bath application to vehicle-treated roots, with 10-minute-recovery intervals between stimulation periods.

C-fibre ADS was quantified by measuring changes in C-fibre latency and width, reflecting the change in average conduction velocity and change in range of the conduction velocities of the C-fibre population, respectively. The latency from the stimulus artefact to the negative peak of the triphasic response was measured, and the change in response latency from the first stimulus was calculated. The change in C-fibre width was analogously calculated, where the width of the C-fibre response was measured from the positive-to-positive peak of the triphasic response. Given that C-fibre ADS is length dependent, the latency/width change was normalised to the length of root stimulated, to negate the influence of varying dorsal root length.

### Patch-clamp electrophysiology

Whole-cell voltage-clamp recordings (holding potential, −70 mV) were made from Fos-EGFP+ spinal neurons from hemizygous Fos-EGFP mice or spinal neurons from WT mice in superficial laminae (lamina I/II). The intracellular solution used in all voltage-clamp recordings contained (in mM): 120 Cs-methanesulfonate, 10 Na-methanesulfonate, 10 ethylene glycol tetraacetic acid (EGTA), 1 CaCl2, 10 4-(2-hydroxyethyl)-1-piperazineethanesulfonic acid (HEPES), 5 2-(triethylamino)-N-(2,6-dimethylphenyl) acetamine chloride (QX-314-Cl), 2 Mg-ATP, pH adjusted to 7.2 with CsOH, osmolarity ∼290mOsm. For current-clamp recording the intracellular solution contained (in mM): 120 K-gluconate, 10 KCl, 0.5 ethylene glycol tetraacetic acid (EGTA), 2 MgCl2, 10 4-(2-hydroxyethyl)-1-piperazineethanesulfonic acid (HEPES), 2.0 Na2-ATP, 0.5 Na-GTP, pH adjusted to 7.2 with KOH, osmolarity, ∼290 mOsm. Junction potential was corrected before recording and 1μM Alexa Fluor 555 hydrazide was included in the recording pipette. Data were recorded and acquired with an Axopatch 200B amplifier and pClamp 10 software (Molecular Devices). Data were filtered at 5kHz and digitised at 10kHz.

Miniature excitatory postsynaptic currents (mEPSCs) were recorded from Fos-EGFP+ neurons in spinal slices obtained from subjects following *in-vivo* intraplantar capsaicin injection (described above) in the presence of 0.5*μ*M TTX, 10*μ*M bicuculline and 1*μ*M strychnine. mEPSCs were recorded at baseline (5 mins), during bath application of TRPV1 agonist, capsaicin (1*μ*M, 5 mins), to pharmacologically potentiate excitatory input,[63] and during a 10 min recovery period. Potentiation was assessed by comparing mEPSCs recorded in the final 2 min of each recording period.

Monosynaptic primary afferent input to spinal neurons was identified as described previously[16; 17; 58; 59]. Appropriate stimulus parameters were verified in CAP recordings from isolated mouse dorsal roots (data not shown, threshold: Aβ range 3-6µA, Aδ range 15-30µA, C range 30-70µA; conduction velocity: Aβ range 0.65-5.54 m/s, Aδ range 0.35-0.7m/s, C range 0.13-0.24m/s) and were consistent with previously reported adult mouse data[6; 13]. Briefly, dorsal roots were stimulated repetitively (X20): Aβ, 10 μA/20 Hz; Aδ, 30 μA/2 Hz; C, 500 μA/1 Hz. Monosynaptic A-fibre inputs were defined based on lack of synaptic failures and a stable latency (≤2 ms). C-fibre inputs were considered monosynaptic if they displayed no synaptic failures, regardless of whether there was latency variability[40]. Evoked EPSCs (eEPSCs) were recorded in response to low-frequency (0.05 Hz) dorsal root stimulation (X3) at 700 μA (0.1 ms stimulus duration) to activate C-fibre inputs. To assess ADS in monosynaptic C-fibre input to spinal neurons, eEPSCs were recorded in response to trains of X16 stimuli at 2 Hz (700μA, 0.1ms duration). The latency of each eEPSC was measured as the time between the stimulus artefact and the onset of the monosynaptic response, and the change in latency from stimulus 1 was calculated to quantify ADS and was normalised to dorsal root length.

In spinal neurons from naïve WT mice and in a subset of Fos-EGFP+ neurons, eEPSC were recorded at 0.05Hz at 700μA following continuous bath application of 1μM capsaicin for 15mins. The amplitude of the monosynaptic C-fibre input was measured before capsaicin application (at 0 min) and at 5, 10 and 15min following bath application. This was used to limit analysis to neurons with capsaicin sensitive and therefore noxious heat responsive monosynaptic C-fibre inputs in WT tissue and enabled confirmation that monosynaptic C-fibre inputs to Fos-EGFP+ neurons were TRPV1 expressing.

Evoked excitatory post-synaptic potentials (eEPSPs) were recorded, following stimulation of the attached dorsal root at two different stimulation distances (to vary levels of ADS) with 10 min intervals between stimulation periods. One electrode was attached near the DRG end of the dorsal root (long) and the other electrode was attached close to the dorsal root entry zone (short). In these ADS length dependency experiments, latency change data was not normalised to dorsal root length.

Monosynaptic C-fibre input-induced action potentials and net-charge were estimated based on the latency of the monosynaptic C-fibre input, defined in voltage-clamp recording.

### Statistical analysis

To compare the effects of chronic MG, sex, and frequency on C-fibre ADS, area under the curve (AUC) for each group was measured and analysed using three-way analysis of variance (ANOVA), followed by Tukey’s multiple comparisons test if an interaction was observed. If one factor did not interact, data were consolidated by this factor to investigate the effects of the interaction between the remaining factors using two-way ANOVA. Appropriate post-hoc multiple comparisons tests were performed only if an interaction between factors was observed. The AUC of the acute MG effect data within each sex, were analysed using one-way ANOVA. Repeated measures (RM) two-way ANOVA was used to assess the effect of MG/vehicle administration and sex upon behavioural measures.

Paired two-tailed t-test was used to compare the number of noxious heat-induced Fos-EGFP+ neurons in the ipsilateral versus contralateral spinal cord. Two-way ANOVA was used to analyse: mEPSC frequency prior to and after capsaicin application; latency change AUC of ADS in monosynaptic C-fibre inputs to and its impact upon noxious heat responsive spinal neurons in both sexes at different stimulation distances; AUC of the activity-dependent changes in eEPSC peak amplitude and net charge in both sexes at different stimulation distances.

Graphad Prism 9.11 software was used for statistical analysis and graph production. All data are presented as mean ± SEM with n representing sample size and N representing number of animals. P<0.05 was used to indicate statistical significance.

### Materials

All chemicals were obtained from Sigma, except from QX-314-Cl (Alomone Labs), thiourea, low melting point agarose, and Alexafluora 488 (Thermos Fisher)

## Results

### Chronic MG alters frequency-dependent C-fibre activity-dependent slowing in a sex-dependent manner

To assess the impact of chronic MG treatment on afferent C-fibre function, population CAPs were recorded from dorsal roots isolated from naïve rats, incubated with vehicle or 100μM MG for 3h. This incubation period and dose of exogenous MG has been established to model the sensory neuron hyperexcitability induced by patient plasma MG levels[10]. Chronic MG treatment did not affect basic electrical properties of C-fibres. There was no significant effect of chronic MG treatment or sex upon C-fibre activation threshold (data not shown; two-way ANOVA: chronic MG, p=0.26, sex, p=0.28, interaction, p=0.72), amplitude (data not shown; two-way ANOVA: chronic MG, p=0.64, sex, p=0.36, interaction, p=0.39) or average conduction velocity (data not shown; two-way ANOVA: chronic MG, p=0.79, sex, p=0.95, interaction, p=0.44).

C-fibre ADS was confirmed to be frequency-dependent with higher frequencies resulting in greater latency (**Figure 1**) and width (**Figure 2**) changes. Chronic MG treatment affected C-fibre latency changes in a sex-dependent manner (**Figure 1E**). In males, chronic MG treatment reduced latency changes compared to vehicle-treated controls, whereas in females it enhanced latency changes.

**Figure 1.**
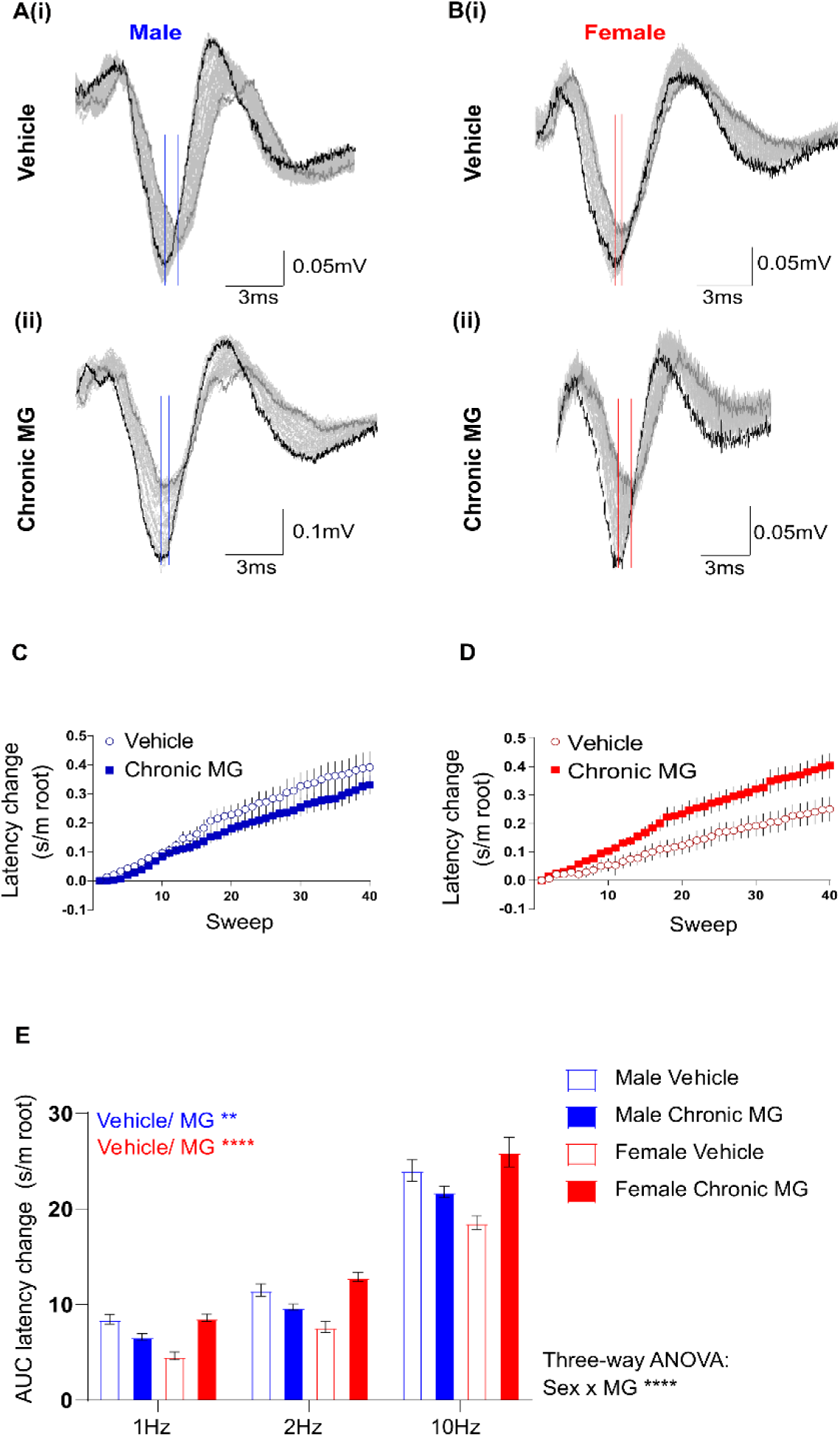
Chronic MG reduces frequency dependent C-fibre latency change in males and enhances it in females. Representative C-fibre CAP recordings, in response to X40 stimuli at 1Hz, from dorsal roots isolated from male (**A**) and female (**B**) naïve rats incubated for 3h in vehicle (**i**) or MG (**ii**). Latency change marked by solid lines (response trace 1 black; 2-39 pale grey; 40 dark grey). Repetitive stimulation at 1Hz results in a progressive C-fibre latency increase in vehicle-treated and MG-treated dorsal roots from males (**C**) and females (**D**). **E**) AUC analysis reveals a significant effect of frequency (****p<0.0001), chronic MG (****p<0.0001), with a sex x chronic MG interaction (****p<0.001) (Three-way ANOVA). Consolidation of the data by frequency to investigate effects of chronic MG within sex showed a statistically significant effect of chronic MG in males (blue font; two-way ANOVA followed by Tukey’s multiple comparisons test, **p=0.008) and females (red font; two-way ANOVA followed by Tukey’s multiple comparisons test, ****p<0.0001) and a statistically significant effect of sex on C-fibre latency change in vehicle and MG-treated dorsal roots (Two-way ANOVA followed by Tukey’s multiple comparisons test, vehicle male vs female, ****p<0.0001 and MG male vs female, ****p<0.0001). Data presented as mean±SEM. Male: vehicle, n=9 (N=6 animals); MG, n=9 (M=5 animals). Female: vehicle, n=10 (N=6), MG, n=9 (N=4 animals).

When C-fibre ADS was assessed as a progressive increase in response width, the impact of chronic MG treatment was also sex-dependent (**Figure 2E**). Chronic MG treatment did not impact width change in males but enhanced it in females, with multiple comparison tests revealing a statistically significant increase in female width change at 2Hz only.

**Figure 2.**
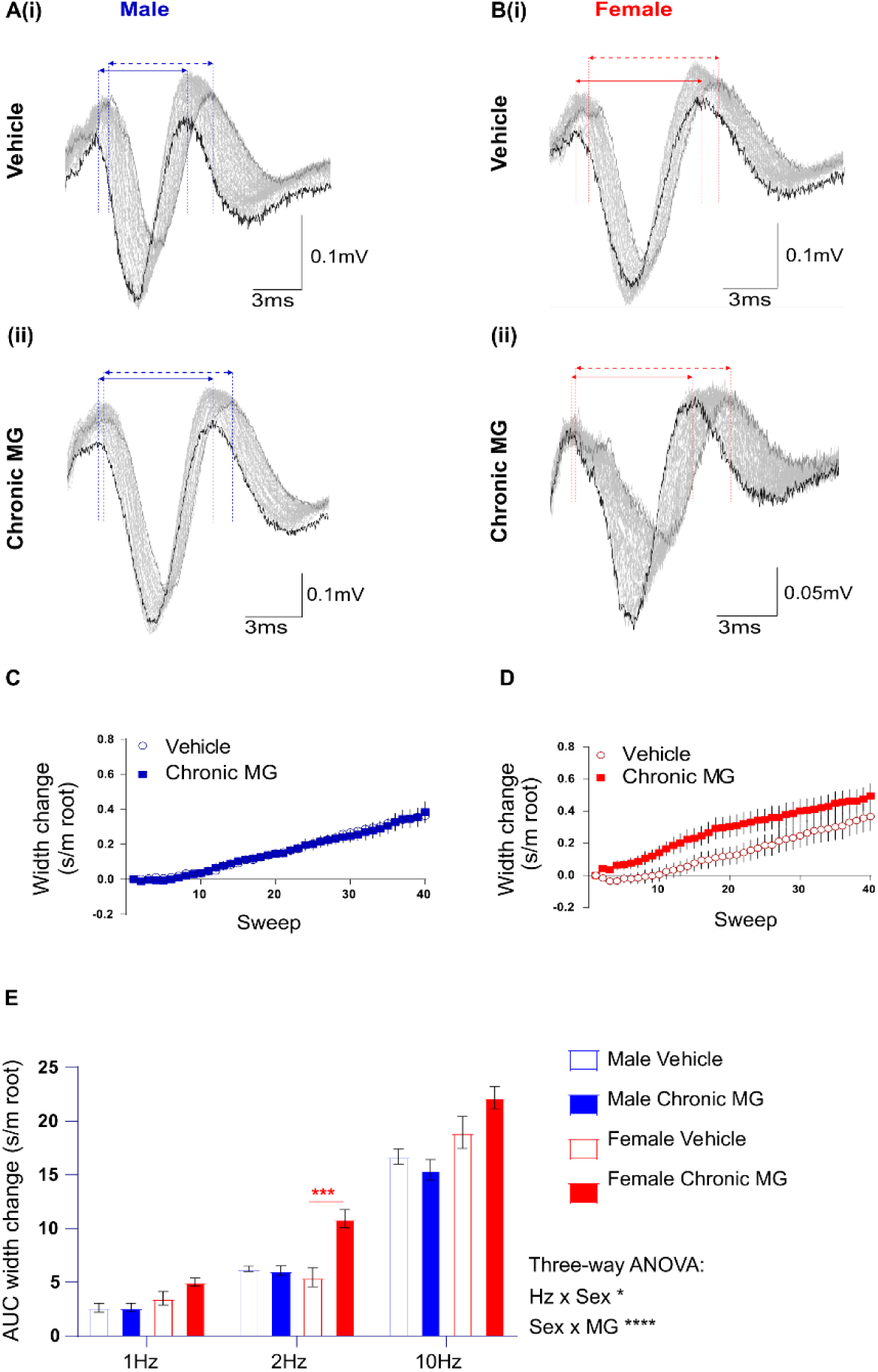
Chronic MG enhances frequency dependent C-fibre width change in females but not in males. Representative C-fibre CAP recordings, in response to X40 stimuli at 2Hz, from dorsal roots isolated from male (**A**) and female (**B**) naïve rats incubated for 3h in vehicle (**i**) or MG (**ii**). Double-headed arrows mark C-fibre response width, the distance from positive-to-positive peaks for the first (solid double-headed arrow) and last (dashed double-headed arrow) of the X40 responses (response trace 1 black; 2-39 pale grey; 40 dark grey). Repetitive stimulation at 2Hz results in a progressive C-fibre width increase in vehicle-and MG-treated dorsal roots from males (**C**) and females (**D**). **E**) AUC analysis reveals a significant effect of frequency (****p<0.0001), sex (****p<0.0001) and chronic MG (**p=0.002), with a frequency x sex interaction (*p=0.026) and a sex x chronic MG interaction (****p<0.0001) (Three-way ANOVA). There was a statistically significant effect of chronic MG in females only (red lines and asterisks) (Three-way ANOVA followed by Tukey’s multiple comparisons test, ***p=0.0002). Data presented as mean±SEM. Male: vehicle, n=9 (N=6 animals), MG, n=9 (M=5 animals). Female: vehicle, n=10 (N=6), MG, n=9 (N=4 animals).

### Acute MG treatment does not impact C-fibre ADS in both sexes

Previous studies have demonstrated that higher MG concentrations can acutely excite nociceptors via TRPA1 activation[4; 18] including along C-fibre axons[19]. To investigate potential involvement of axonal TRPA1 expression, the acute effects of MG (100μM and 1mM) were also studied (**Figure 3**).

**Figure 3.**
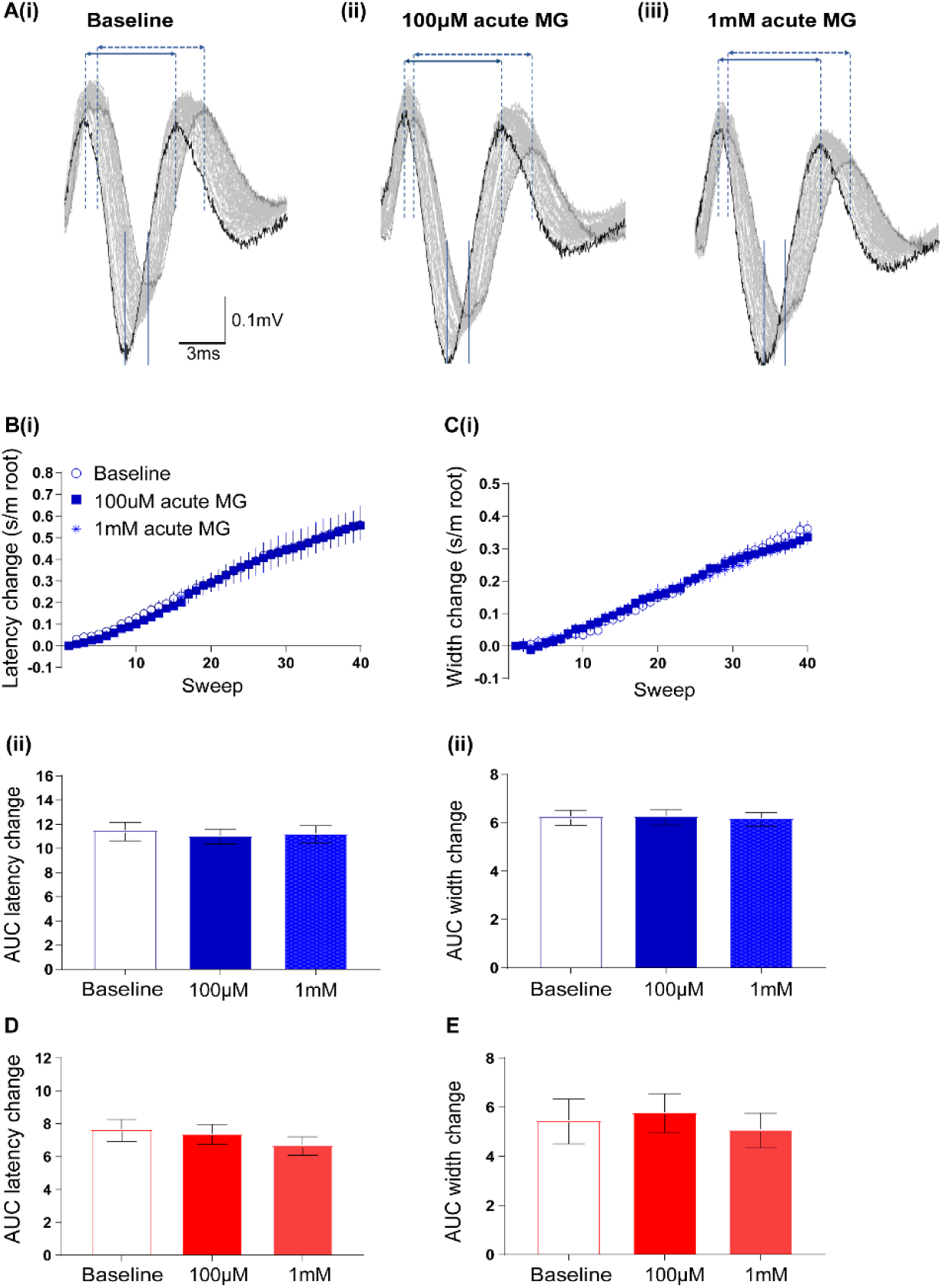
Acute MG treatment has no effect on C-fibre ADS in both sexes. **A)** Representative C-fibre CAP recordings, in response to X40 stimuli at 2Hz, from dorsal roots isolated from naïve male rats at baseline (**i**) and following acute (10min bath) 100μM (**ii**) and 1mM (**iii**) MG application. Latency change marked by solid lines; double-headed arrows mark C-fibre response width, the distance from positive-to-positive peaks for the first (solid double-headed arrow) and last (dashed double-headed arrow lines) of the X40 responses (response trace 1 black; 2-39 pale grey; 40 dark grey). Repetitive stimulation results in a similar levels of progressive C-fibre latency **B**(**i**) and width **C**(**i**) increase at baseline and following bath application of 100μM and 1mM MG. AUC analysis of C-fibre latency change in males (**Bii**; One-way ANOVA: p=0.865) and females (**D**; One-way ANOVA: p=0.469) and width change in males (**Cii**; One-way ANOVA: p=0.962) and females (**E**; One-way ANOVA: p=0.808) reveals no significant effect of acute MG treatment. Data presented as mean±SEM. Male: n=9 (N=6 animals); Female: n=10 (N=6).

Acute MG treatment did not alter male C-fibre latency (**Figure 3B**) or width (**Figure 3C**) changes and similarly did not alter female C-fibre latency (**Figure 3D**) or width (**Figure 3E**) changes. Therefore, it is unlikely that the chronic MG sex-dependent changes in C-fibre ADS involve axonal TRPA1.

### Systemic MG induced heat hyperalgesia in males but not females

Systemic administration of MG, to mimic patient plasma levels, induces heat hyperalgesia in male mice[10]. Similarly, systemic administration of MG reduced the withdrawal latency to a noxious heat stimulus, 3 hours post MG injection, in juvenile male rats (**Figure 4A)**. Interestingly, systemic MG administration did not alter the withdrawal latency to a noxious heat stimulus in juvenile female rats (**Figure 4B**). To directly compare with the previous findings in adult male mice[10], data were also presented as a change from baseline measures, which confirmed the sex-dependent development of heat hyperalgesia in males only (**Figure 4C**). Systemic MG treatment did not alter the withdrawal threshold to a punctate mechanical stimulus in juvenile rats of both sexes, when assessed 3 hours post-injection (Data not shown; Two-way ANOVA: MG, p=0.273, sex, p=0.727, MG x sex interaction, *p*=0.770).

**Figure 4.**
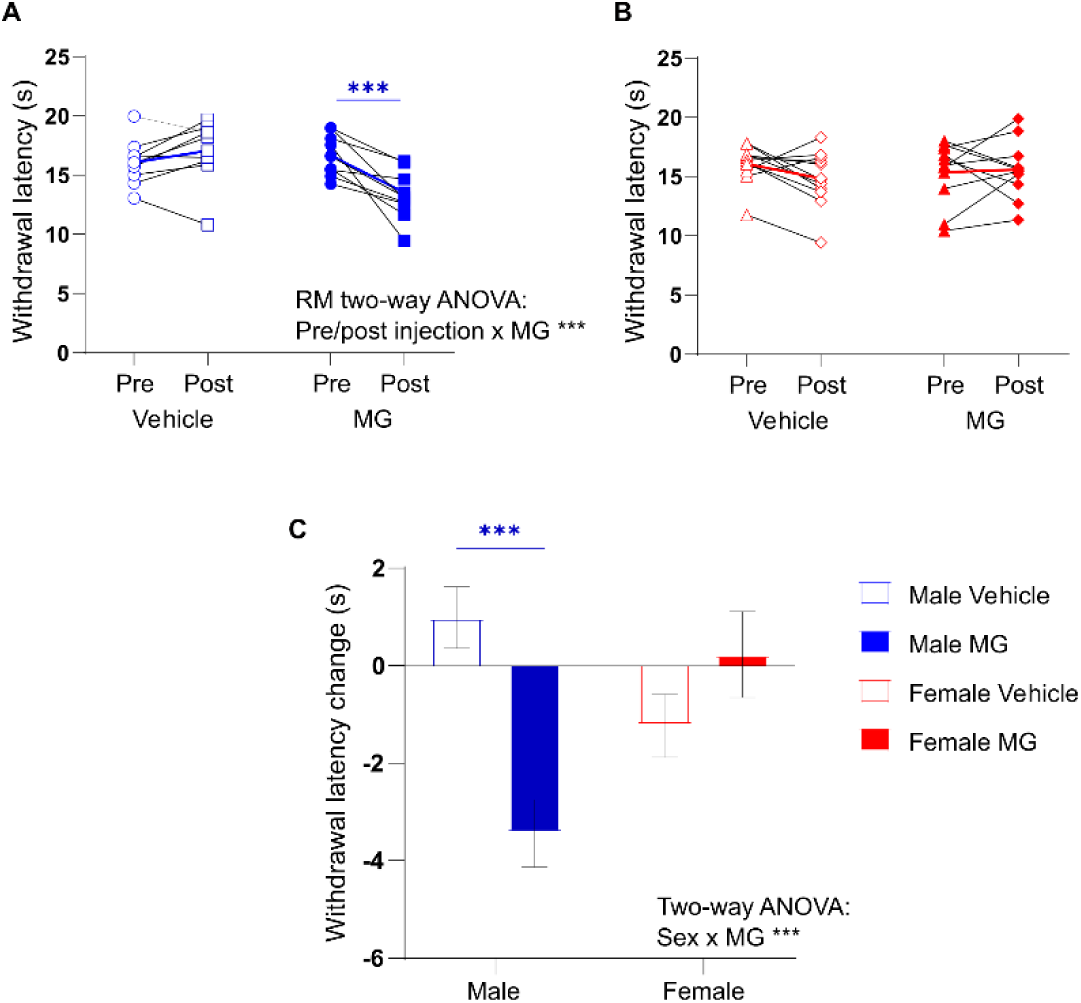
Systemic MG administration induces heat hyperalgesia in males but not in females. Hindpaw withdrawal latency to a radiant noxious heat stimulus (Hargreaves) pre- and post-i.p injections with vehicle or 5μg MG in male (**A**) and female (**B**) juvenile rats. Systemic administration of MG decreased the withdrawal latency to a noxious heat stimulus in males (RM Two-way ANOVA: vehicle vs MG, p=0.064, pre-vs post-injection, p=0.019, pre/post injection x MG interaction, ***p=0.0002 followed by Sidak’s multiple comparisons test, pre-vs post-MG, ***p=0.0001), but not in females (Two-way ANOVA: vehicle vs MG, *p*=0.987, pre-vs post-injection, p=0.367, pre/post injection x MG interaction, p=0.191). Data are presented per individual animal with thicker blue and red lines representing the mean values for males and females, respectively. **C**) Hindpaw withdrawal latency to a radiant noxious heat stimulus, following MG or vehicle administration, presented as a change from baseline latency reveals a significant latency reduction following MG in males but not in females (Two-way ANOVA: MG, *p=0.048, sex, p=0.324, interaction, ***p=0.0002 by followed by Sidak’s multiple comparisons test, male vehicle vs male MG, *\*\*\**p=0.0009, female vehicle vs female MG, *p*=0.625). Data presented as mean±SEM. Male: vehicle, n=9, MG, n=10. Female: vehicle, n=11, MG, n=10.

### Impact of ADS on the processing of monosynaptic C-fibre inputs by noxious heat-responsive spinal neurons

Our prior work, in *ex vivo* spinal network recordings, suggests that ADS regulates the spinal processing of C-fibre inputs [16]. Given the present finding that chronic MG regulates both C-fibre ADS and behavioural heat sensitivity in a sex-dependent manner we hypothesised that ADS regulates the processing of monosynaptic C-fibre inputs by noxious heat responsive spinal neurons. To explore this, we utilised activity-marker transgenic mice, specifically Fos-EGFP mice[8] to enable patch-clamp recording from noxious heat responsive neurons in spinal slices.

We first validated the approach to show that spinal neurons with Fos-EGFP expression, induced by prior *in vivo* intra-plantar capsaicin injection, have capsaicin sensitive inputs. Indeed, Fos-EGFP+ dorsal horn neurons (**Figure 5Ai-ii**), displayed a significant increase in mEPSC frequency in response to bath applied capsaicin (**Figure 5B, C**), whereas nearby Fos-EGFP negative neurons showed no change in mEPSC frequency (**Figure 5C**).

**Figure 5.**
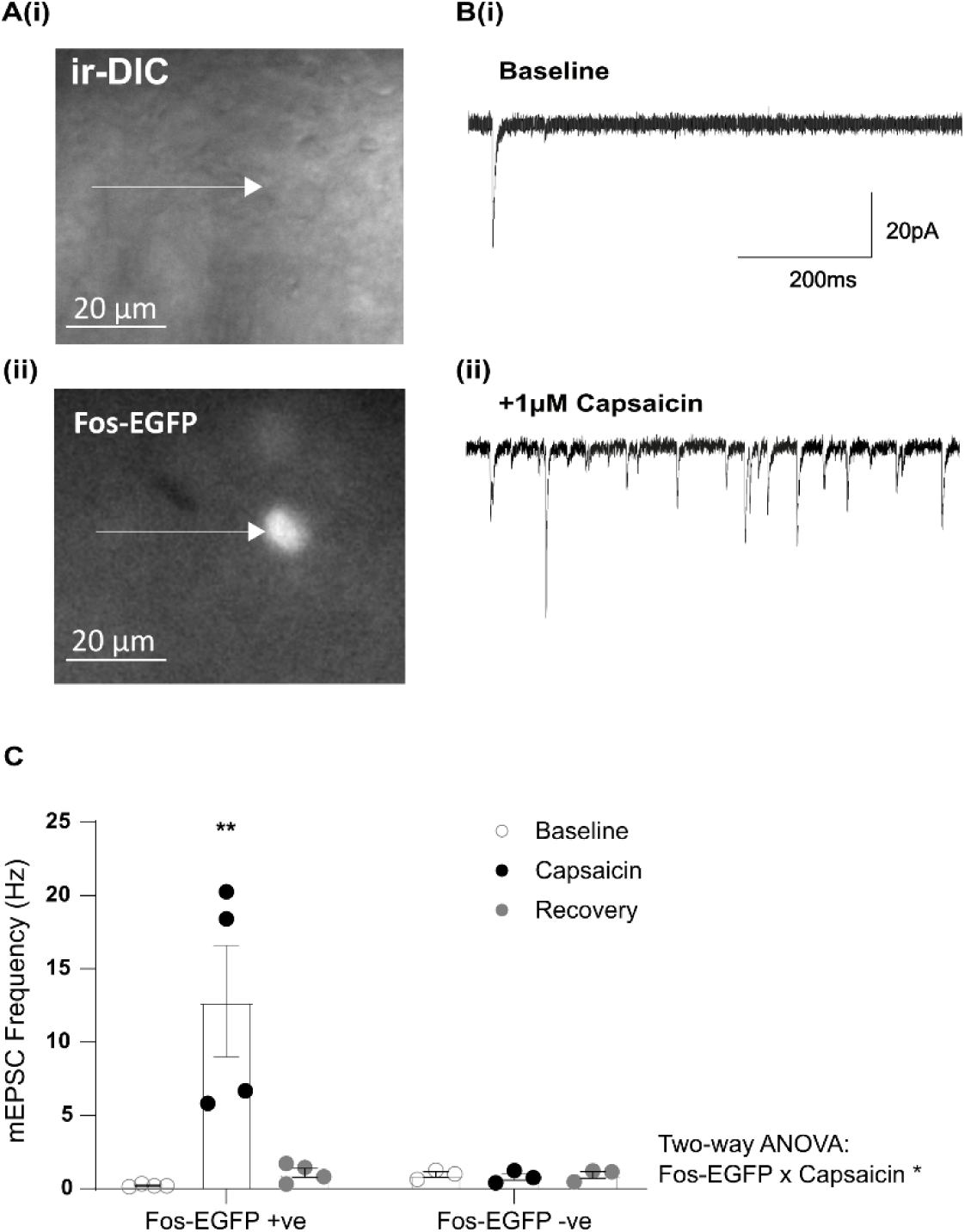
In vivo intra-plantar capsaicin injection leads to Fos-EGFP expression in spinal neurons with capsaicin responsive inputs. **A)** Example image of Fos-EGFP expressing neuron, induced by intraplantar capsaicin injection 2h prior, in an *ex vivo* spinal cord slice preparation, visualised with ir-DIC (**i**) and fluorescence (**ii**) microscopy. **B**) Representative mEPSC traces recorded at baseline (**i**) and during bath application of 1μM capsaicin (**ii**). **C**) Analysis of the mEPSC frequency at baseline, during and after wash out (recovery) of capsaicin in Fos-EGFP+ neurons and in nearby Fos-EGFP negative neurons revealed an effect of capsaicin that is dependent on cell type (*p=0.01), with capsaicin increasing mEPSC frequency in Fos-EGFP+ neurons but not in nearby Fos-EGFP negative neurons (Two-way ANOVA followed by Sidak’s multiple comparisons test, **p=0.001, baseline vs capsaicin; **p=0.002, capsaicin vs recovery). Data presented as mean±SEM. Fos-EGFP positive cells, n=4 (N=4); Fos-EGFP negative cells, n=3 (N=3).

*In vivo* noxious heat stimulation also led to Fos-EGFP expression in spinal dorsal horn neurons (**Figure 6Ai**) with significantly more Fos-EGFP+ neurons ipsilateral, compared with contralateral, to the applied stimulus (**Figure 6Aii**). Voltage-clamp recording demonstrated that they received monosynaptic C-fibre input, which exhibited ADS (**Figure 6Bi-ii**). To test whether the noxious heat-induced Fos-EGFP+ neurons have monosynaptic C-fibre inputs that are capsaicin-sensitive, i.e. TRPV1 expressing as would be predicted[12], eEPSCs were recorded prior to and during the continued presence of 1μM capsaicin. Capsaicin application led to a progressive reduction in the peak amplitude of the monosynaptic C-fibre eEPSCs (**Figure 6C-D**).

**Figure 6.**
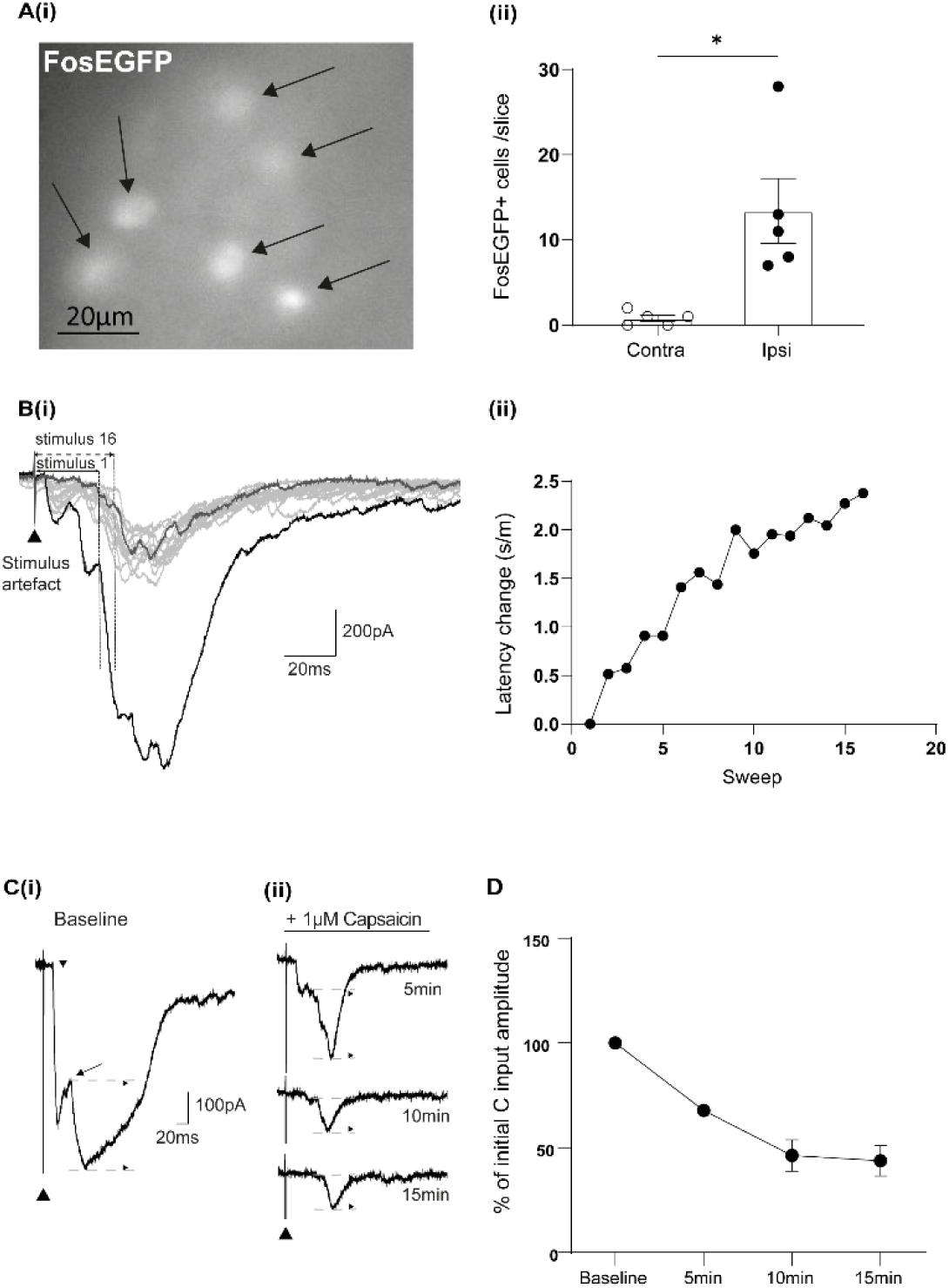
Natural noxious heat stimulation induces Fos-EGFP expression in spinal neurons, including neurons with monosynaptic C-fibre input that display ADS. **A)** Noxious heat stimulation of the hindpaw by brief immersion in a 52°C water bath induced Fos-EGFP expression in neurons (marked by arrows) in the superficial dorsal horn of lumbar (L4/5) spinal cord (**i**). There were significantly more Fos-EGFP+ neurons in the superficial dorsal horn of the spinal cord, ipsilateral compared with contralateral to the stimulus (**Aii**; Paired two tailed t-test, *p=0.027). Data presented as mean±SEM, n=5 slices (N=2). **Bi**) Representative monosynaptic C-fibre eEPSC recording from a Fos-EGFP+ spinal neuron following repetitive stimulation (X16 stimuli at 700μA at 2Hz). Response trace 1 black; 2-15 pale grey; 16 dark grey. Monosynaptic C-fibre eEPSC response latency to the first (solid double-headed arrow) and 16^th^ stimuli (dashed double-headed arrow) is shown (vertical lines indicate onset of monosynaptic C-fibre component). **Bii**) Repetitive stimulation results in a progressive latency increase demonstrating the ADS phenomenon in a monosynaptic C-fibre input to a Fos-EGFP+ noxious heat responsive spinal neuron. **C**) Representative eEPSC recordings (X3 average) at 500μA at 0.05Hz showing A-fibre eEPSC latency (grey arrow) and monosynaptic C-fibre eEPSC latency (black arrow) and monosynaptic C-fibre amplitude (vertical double-headed arrow) at baseline (**i**) and 5min, 10min and 15min following bath application of 1μM capsaicin (**ii**). **D**) C-fibre eEPSC peak amplitude as a percentage of the initial C-fibre eEPSC peak amplitude at different time points following capsaicin bath application. Data presented as mean±SEM, n=4 cells (N=4).

The length dependency of ADS[49; 64] was used to investigate the impact of different levels of ADS on the processing of monosynaptic C-fibre inputs by noxious heat responsive spinal neurons.

Repetitive stimulation resulted in a clear ADS in monosynaptic C-fibre inputs at both short and long stimulation distances, reflected by the progressive increase in the C-fibre eEPSC latency (**Figure 7A**). Shorter stimulation distances resulted in a significantly less pronounced latency increase in monosynaptic C-fibre eEPSCs, with no significant difference between males and females (**Figure 7B**). In addition, there was no significant effect of sex upon activation threshold (unpaired two-tailed t-test: p=0.243), conduction velocity (unpaired two-tailed t-test: (p>0.999) or initial latency (unpaired two-tailed t-test: p=0.949) of the monosynaptic C-fibre inputs to the noxious heat responsive spinal neurons (data not shown).

**Figure 7.**
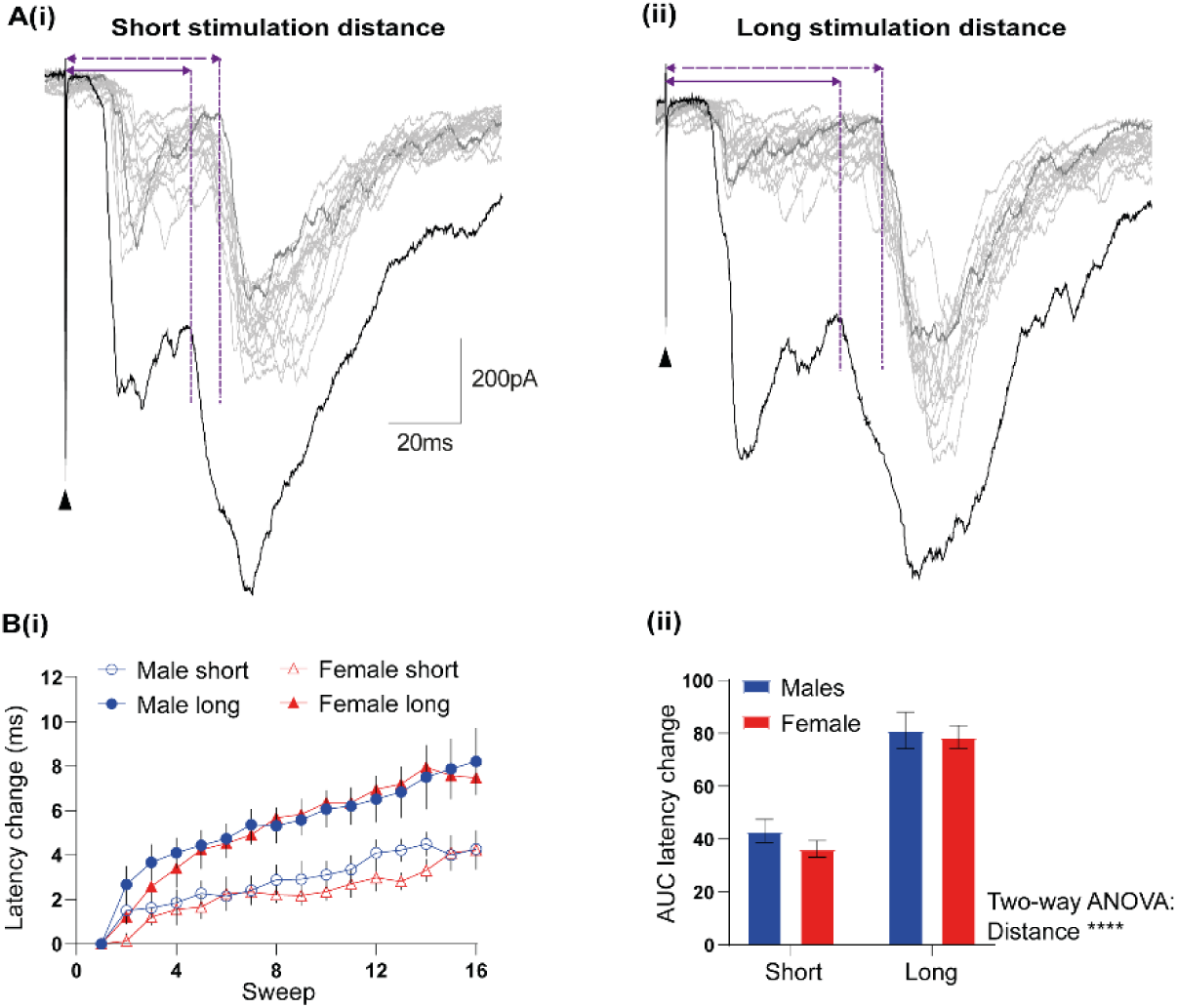
Length-dependent modulation of progressive latency change ‘ADS’ in monosynaptic C-fibre input to noxious heat responsive spinal neurons. **A)** Representative eEPSC recordings of monosynaptic C-fibre input in response to X16 stimuli at 2Hz at 700μA at short (**i**) and long (**ii**) stimulation distance (response trace 1 black; 2-15 pale grey; 16 dark grey). Monosynaptic C-fibre eEPSC response latency to the first (solid double-headed arrow) and the 16^th^ (dashed double-headed arrow) stimulus is shown (vertical dotted lines indicate onset of monosynaptic C-fibre component). **B**) Progressive increase in response latency at short and long stimulation distances in both sexes during repetitive stimulation of monosynaptic C-fibre input to Fos-EGFP+ noxious heat responsive spinal neurons (**i**). AUC analysis (**ii**) revealed a significant effect of stimulation distance (****p<0.0001), with more pronounced ADS at long stimulation distances in both sexes (p=0.371) and no interaction between the two factors (p=0.675) (Two-way ANOVA). Data presented as mean±SEM. Males, n=7 (6x Fos-EGFP+ and 1x WT capsaicin-sensitive) (N=5); females, n=6 (5x Fos-EGFP+ and 1x WT capsaicin-sensitive) (N=4).

To assess the impact of the degree of ADS in monosynaptic C-fibre inputs upon the activity of noxious heat responsive spinal neurons, eEPSP recordings were conducted in Fos-EGFP+ noxious heat responsive neurons from spinal cord slices with an attached dorsal root stimulated at a short and long distance in both sexes (**Figure 8**). Given dorsal root repetitive stimulation induces spinal summation in ex vivo network recordings that is enhanced at shorter distance (reduced ADS) stimulation[16] it was predicted that shorter distance repetitive stimulation would enhance summation of action potential firing in Fos-EGFP+ neurons. To enable comparison of distance and sex, data were normalised to the number of action potentials induced by the first stimulus in a given condition to account for any potential differences in initial action potential number between conditions. Repetitive stimulation was observed to progressively reduce, rather than summate, the normalised number of C-fibre-evoked action potentials at short and long stimulation distances in both sexes (**Figure 8C**). Nevertheless, the progressive decrease in C-fibre-evoked action potential firing was indeed length dependent; greater at short compared to long stimulation distances in both sexes.

**Figure 8.**
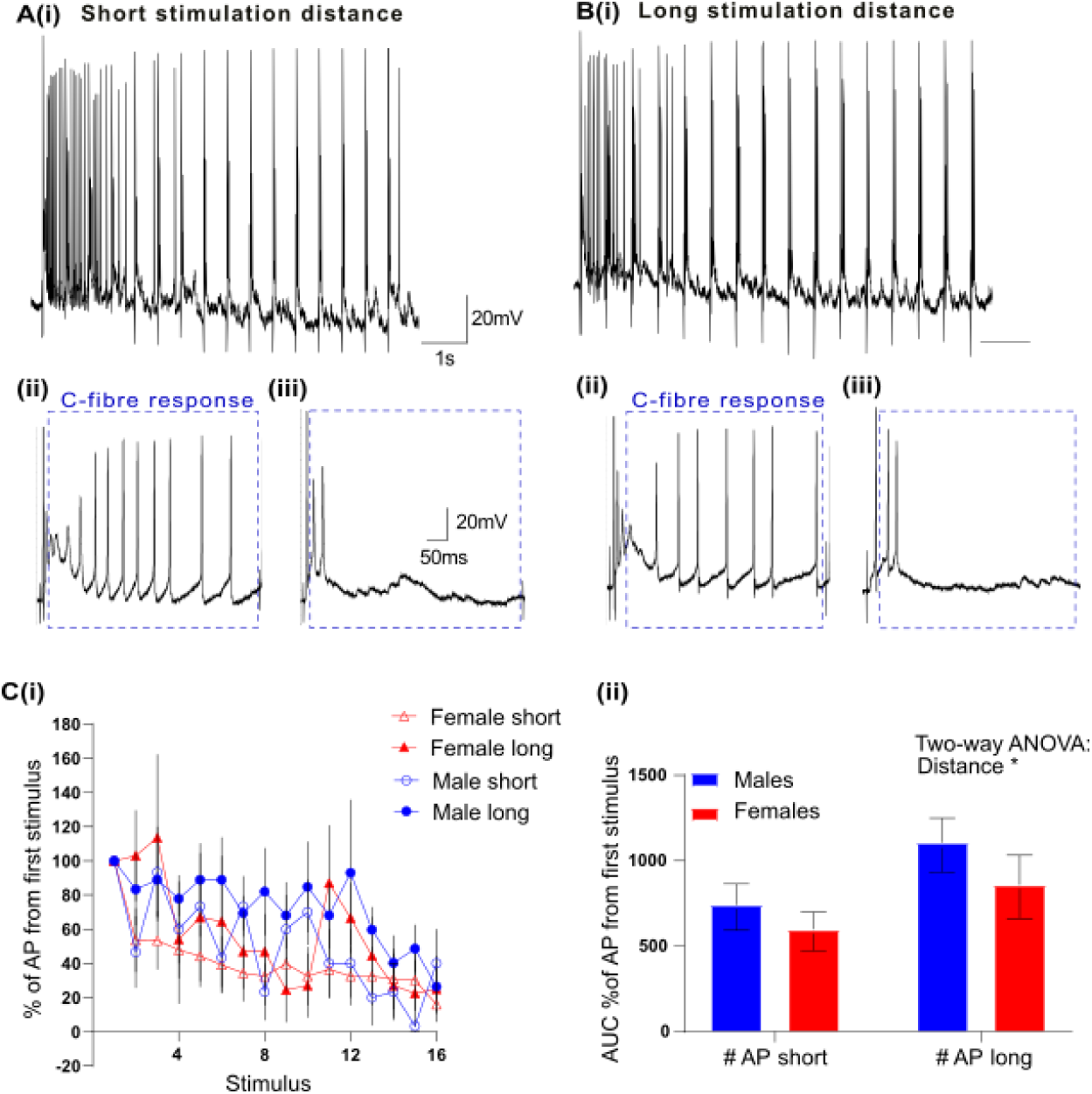
Length-dependent modulation of repetitive C-fibre evoked action potential firing in noxious heat responsive neurons. Representative eEPSP recordings from Fos-EGFP+ heat responsive neurons in spinal cord slices with attached dorsal roots, repetitively stimulated (X16 stimuli at 2Hz at 700μA) at a short (**A**) and at a long (**B**) dorsal root distance. An expanded timescale shows the number of action potentials evoked at the first (**ii**) and the 16^th^ (**iii**) stimulus. C-fibre –evoked action potentials are marked by the dashed boxes. **C**) Progressive decrease in the normalised number of action potentials at short and long stimulation distances in both sexes (**i**) following repetitive stimulation of monosynaptic C-fibre input to Fos-EGFP+ heat responsive spinal neurons. AUC analysis (**ii**) showed significantly more pronounced action potential firing at long compared to short stimulation distances (*p=0.038) in both sexes (p=0.178), with no interaction between the two factors (p=0.71) (Two-way ANOVA). Data presented as mean±SEM. Males, n=6 (N=4); females, n=5 (N=2).

To assess whether the observed impact of ADS upon the activity of noxious heat responsive spinal neurons could be explained by timing-dependent presynaptic regulation of monosynaptic C-fibre inputs (eEPSC peak amplitude) or in timing-dependent polysynaptic C-fibre evoked activity (net charge) these were quantified. There was no significant effect of distance or sex on the normalised peak amplitude with repetitive stimulation (**Figure 9A**). Similarly, there was no significant effect of distance or sex on normalised net charge with repetitive stimulation (**Figure 9B**).

**Figure 9.**
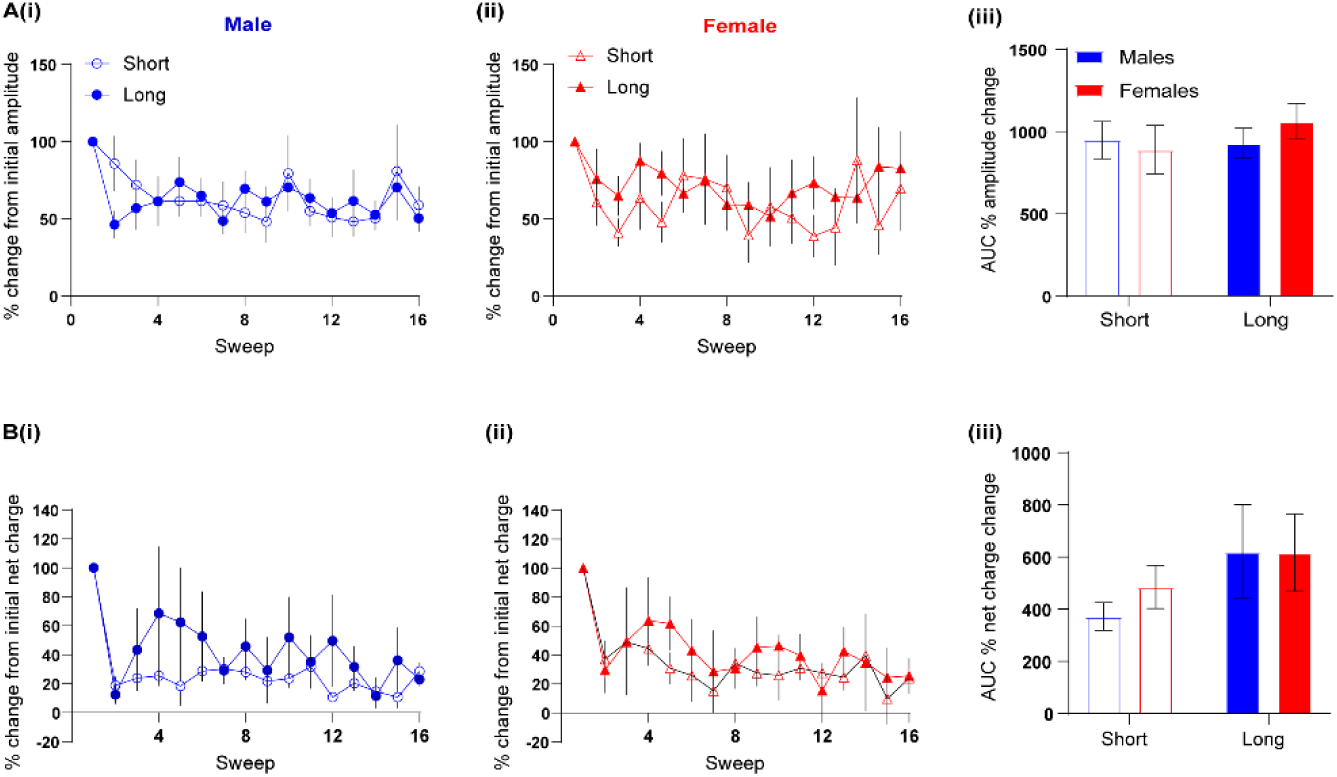
Stimulation distance does not impact normalised C-fibre eEPSC peak amplitude or net charge following repetitive stimulation. Normalised peak amplitude (**A**) and net charge (**B**) per stimulus at short and long stimulation distances in males (**i**) and females (**ii**) following repetitive stimulation of monosynaptic C-fibre input to heat responsive spinal neurons. AUC analysis reveals no significant differences in normalised peak amplitude (**Aiii**; Two-way ANOVA: stimulation distance, p=0.505, sex, p=0.756, stimulation distance x sex interaction, p=0.418) or in the net charge (**Biii**; Two-way ANOVA: stimulation distance, p=0.150, sex, p=0.673, stimulation distance x sex interaction, p=0.654). Data presented as mean±SEM. Males, n=7 (6x Fos-EGFP and 1x WT capsaicin-sensitive) (N=5); females, n=6 (5x Fos-EGFP+ and 1x WT capsaicin-sensitive) (N=4).

## Discussion

Chronic MG treatment, in isolated rat dorsal roots, resulted in sex-dependent modulation of C-fibre ADS; it was reduced in males but increased in females. Systemic administration of MG induced heat hyperalgesia in male but not female rats. Experimental manipulation of ADS showed that it can influence action potential firing in noxious heat responsive spinal neurons in Fos-EGFP mice. We propose that MG sex-dependent regulation of ADS, by influencing spinal processing of noxious heat inputs may contribute to the sex-dependent impact of MG on heat pain sensitivity.

### MG-sex-dependent modulation of C-fibre ADS

Chronic MG, that can modulate Na_V_1.7 and Na_V_1.8[10], regulates C-fibre ADS in a sex dependent manner. This is consistent with C-fibre ADS, that is influenced by the ratio of Na_V_1.7 to Na_V_1.8[45], also being sex-dependent[16; 60]. Collectively these findings suggest sex differences in the functional expression of Na_V_1.7 and Na_V_1.8 along C-fibre axons that is supported by low-dose TTX sex-dependent modulation of C-fibre ADS[60].

At the level of the DRG cell body there is a lack of evidence for sex-dependent expression of Na_V_1.7 and Na_V_1.8[36; 37] or sex differences in Na_V_ biophysical properties[23]. However, a lack of sex difference at the cell body level does not preclude sex differences in axonal Na_V_1.7 and Na_V_1.8 function. There is evidence for sex-specific CRMP2-SUMOylation dependent trafficking of Nav1.7[39]. In addition, as observed for other nociceptor proteins, there could be sex-dependent hormonal regulation and translation[43; 44].

Given the present findings it would be of interest to compare C-fibre ADS profiles in both sexes in preclinical and clinical diabetic neuropathy studies, but to date, studies are male only[21; 61] or not separated by sex[33], respectively.

### MG and painful diabetic neuropathy

The chronic MG reduced C-fibre ADS profile in males, should facilitate spinal pain processing[16], consistent with the observed systemic MG induction of heat hyperalgesia in males. This contrasts the chronic MG enhanced C-fibre ADS in females and lack of induction of heat hyperalgesia with systemic MG. However, in the clinic it is female sex that is a significant risk factor for painful diabetic neuropathy[2; 11] and females report more severe pain symptoms[2]. However, the directionality of sex differences in pain depends on genotype[38]. Therefore, it would be interesting to know whether C-fibre ADS in humans displays a sex difference and its directionality, given that sex-dependent C-fibre ADS was predictive of a sex-dependent action of MG.

MG also induces pain symptoms via other mechanisms. Low dose MG (∼1µm) can activate the integrated stress response (ISR) to activate IB4-positive nociceptors and drive MG induced mechanical hypersensitivity[7]. However, given the lack of systemic MG-induced mechanical hypersensitivity in the behavioural assays, ISR is an unlikely contributor to the present findings. Higher (mM) MG levels, exceeding that used in the present study, can activate nociceptors via TRPA1 activation and directly induce pain responses[4; 18]. Moreover, elevated MG levels promote oxidative stress and mitotoxicity[14; 48]; well established pathogenic mechanisms that contribute to neuropathic pain[9]. It should be noted that an association between methylglyoxal and painful diabetic neuropathy has been evidenced in some[10] but not all studies[26] but this may reflect the shorter diabetes duration in the latter study.

### ADS regulation of spinal noxious heat processing

Our prior work showed that reduced ADS (shorter stimulation length) enhanced spinal summation in network recordings[16]. In this study, individual noxious heat responsive spinal neurons displayed a decline in action potential firing with repeated stimulation rather than summation. This was greater at shorter stimulation distances, consistent with C-fibre ADS regulation of spinal neuron activity. This may reflects a timing-dependent intrinsic property that influences action potential firing, such as calcium-activated potassium currents that remains to be explored[46].

The lack of summation at the individual neuron level is consistent with the observation, that although evident[24], summation is not frequently observed in superficial dorsal horn neurons[25; 51]. Interestingly, the observed decline in firing, during a stimulus train, is akin to the ‘declining’ calcium response induced by noxious heat in inhibitory neurons as compared with the ‘sustained’ calcium response observed in excitatory neurons[55]. The use of Fos-EGFP transgenic mice to identify noxious heat responsive neurons may have biased sampling to inhibitory neurons[35] which are more likely to receive monosynaptic C-fibre input[27; 28; 50]. A more dramatic decline in action potential firing with repetitive stimulation in inhibitory neurons, resulting from reduced C-fibre ADS, would presumably enhance summation at a spinal network level. Therefore, the chronic MG reduced C-fibre ADS profile in males is consistent with MG-induced heat hyperalgesia in males. Future studies, should specifically compare C-fibre ADS regulation of activity in inhibitory vs excitatory subpopulations which could further understanding of sex differences in spinal pain processing[15].

## Acknowledgements

The work was supported by a Biomedical Sciences International Career development PhD Studentship, Zhejiang University and The University of Edinburgh and a Carnegie Trust for the Universities of Scotland vacation scholarship. Staff of the Bioresearch and Veterinary Services at the University of Edinburgh. The authors have no conflicts of interest.

